# XA4C: eXplainable representation learning via Autoencoders revealing Critical genes

**DOI:** 10.1101/2023.07.16.549209

**Authors:** Qing Li, Yang Yu, Pathum Kossinna, Theodore Lun, Wenyuan Liao, Qingrun Zhang

## Abstract

Machine Learning models have been frequently used in transcriptome analyses. Particularly, Representation Learning (RL), e.g., autoencoders, are effective in learning critical representations in noisy data. However, learned representations, e.g., the “latent variables” in an autoencoder, are difficult to interpret, not to mention prioritizing essential genes for functional follow-up. In contrast, in traditional analyses, one may identify important genes such as Differentially Expressed (DiffEx), Differentially Co-Expressed (DiffCoEx), and Hub genes. Intuitively, the complex gene-gene interactions may be beyond the capture of marginal effects (DiffEx) or correlations (DiffCoEx and Hub), indicating the need of powerful RL models. However, the lack of interpretability and individual target genes is an obstacle for RL’s broad use in practice. To facilitate interpretable analysis and gene-identification using RL, we propose “Critical genes”, defined as genes that contribute highly to learned representations (e.g., latent variables in an autoencoder). As a proof-of-concept, supported by eXplainable Artificial Intelligence (XAI), we implemented eXplainable Autoencoder for Critical genes (XA4C) that quantifies each gene’s contribution to latent variables, based on which Critical genes are prioritized. Applying XA4C to gene expression data in six cancers showed that Critical genes capture essential pathways underlying cancers. Remarkably, *Critical genes has little overlap with Hub or DiffEx genes, however, has a higher enrichment in a comprehensive disease gene database (DisGeNET), evidencing its potential to disclose massive unknown biology*. As an example, we discovered five Critical genes sitting in the center of Lysine degradation (hsa00310) pathway, displaying distinct interaction patterns in tumor and normal tissues. In conclusion, XA4C facilitates explainable analysis using RL and Critical genes discovered by explainable RL empowers the study of complex interactions.

**Author Summary:** We propose a gene expression data analysis tool, XA4C, which builds an eXplainable Autoencoder to reveal Critical genes. XA4C disentangles the black box of the neural network of an autoencoder by providing each gene’s contribution to the latent variables in the autoencoder. Next, a gene’s ability to contribute to the latent variables is used to define the importance of this gene, based on which XA4C prioritizes “Critical genes”. Notably, we discovered that Critical genes enjoy two properties: (1) Their overlap with traditional differentially expressed genes and hub genes are poor, suggesting that they indeed brought novel insights into transcriptome data that cannot be captured by traditional analysis. (2) The enrichment of Critical genes in a comprehensive disease gene database (DisGeNET) is higher than differentially expressed or hub genes, evidencing their strong relevance to disease pathology. Therefore, we conclude that XA4C can reveal an additional landscape of gene expression data.

## INTRODUCTION

Machine learning (ML) models play increasingly important roles in gene expression analyses. Among many ML techniques, representation learning (RL) has the potential to bring a breakthrough, due to its ability to deconvolute nonlinear structures and denoise confounders [1]. For example, in the data preprocessing stage, in contrast to traditional statistical models that remove principal components to adjust data [2], modern tools can run RL to eliminate noise and possibly non-linear structures in the data [3-7]. Autoencoders (AE) have been extensively utilized to develop various tools for processing expression data [3-5, 8]. A notable feature of AEs is their ability to learn the hidden representations of input data despite the input being noisy and heterogeneous, leading to “latent variables” that are cleaner and more orthogonal for next stages of analysis.

However, such learned representations, although enjoying desirable statistical properties, are difficult to interpret. Therefore, in practice, researchers and clinical practitioners are left with manual inspection of data to decide whether to conduct experimental follow-up or clinical investigations. Additionally, traditional expression analyses naturally provide a list of prioritized genes. For instances, one may identify genes of importance such as Differentially Expressed genes (DiffEx) based on individual genes’ marginal effects [9-11] and Differentially Co-Expressed genes [12] based on genes’ pairwise correlations [13-15]. Also, Hub genes can be prioritized based on the connectivity of the nodes in a gene-gene co-expression network [13]. In contrast, the (usually uninterpretable) learned representations do not offer analogues for the experimentalists or clinicians to follow up. Although several tools supporting interpretable ML models are available [4, 8, 16], they do not offer individual candidate genes based on learned representations (**Discussion**). Therefore, to facilitate broader use of RL of expression data, interpretable tools that prioritize candidate genes are urgently needed.

Herein, supported by state-of-the-art development in eXplainable Artificial Intelligence (XAI) [17], we developed a tool, XA4C (eXplainable Autoencoder for Critical genes), to facilitate explainable analysis and prioritization of individual genes. Technically, XA4C is composed of two main components: First XA4C offers optimized autoencoders to process gene expressions at two levels: whole transcriptome (global) autoencoder, and single pathway (local) autoencoders (**Materials & Methods**). Second, using SHapley Additive exPlanations (SHAP) [18-20], a pioneering method inspired by the popular economic concept of “Shapley Value” quantifying the contribution of a player in a game, XA4C quantifies individual gene’s contribution to the learned latent variables in an autoencoder (**Materials & Methods**), and aggregate them to form “Critical index” for each gene (**Materials & Methods**). These Critical indexes will be used to prioritize Critical genes based on user specified cutoff, e.g., 1%.

The term “*Critical gene*”, reflecting genes substantially explaining of latent variables is comparable to the popular term “Hub gene” which is defined as the genes with high connectivity in a co-expression network [13]. These two could be considered in parallel because “Critical genes” and “Hub genes” both play a sensible role in gene-gene interactions, although in different forms of representations in an interaction network. In other words, “Hub genes” contribute to the surrounding genes in the co-expression network through correlations in the plain representation, whereas “Critical genes” contribute to linked genes through the latent variables in an autoencoder through explaining their variations. As hidden states presumably incorporate complex correlation structures, the links between genes and hidden states may be considered the analogue of links in a co-expression network, hence the analogous relationship between Hub genes and Critical genes.

By applying XA4C to cancer data offered by The Cancer Genome Atlas [21], or TCGA (**Materials & Methods**), we revealed sensible genes and pathways through SHAP-based explanations. We also carried out thorough investigations of Critical Genes by generating summary statistics in comparison to other conventional means, i.e., DiffEx genes and Hub genes. We observed important properties of Critical genes: first, the overlaps between Critical genes with DiffEx or Hub genes are quite low; and second, Critical genes’ enrichment in a comprehensive disease-gene database is higher than the ones of DiffEx and Hub genes. <u>These</u> <u>indicate that Critical genes indeed revealed new candidates into the pathology, and they are</u> <u>even more sensible than the DiffEx and Hub genes revealed by traditional analysis</u>. As a step further, we analyzed the data and discovered Critical genes (that are not DiffEx or Hub) altering the interaction patterns in Lysine degradation (hsa00310) pathway.

## RESULTS

### The XA4C model

Autoencoder (AE) is a RL model that learns representations of input datasets in an unsupervised way [1, 22]. Based on an AE with fully connected neural network, XA4C first learns representations of gene expressions (**Materials & Methods**; **Fig 1A)**. Briefly, input gene expression profiles are passed through the encoder network to learn low-dimensional representations through a bottleneck. The decoder is symmetrical to the encoder counterpart to recover the gene expressions. The loss function of training the parameters in the AE is Mean Squared Error (MSE) between input and output, the default for continuous variables. To quantify each gene’s contribution to the latent variables, XA4C employs eXtreme Gradient Boosting (XGBoost) [23], an ensemble tree model [24] between the input genes and latent variables (**Materials & Methods**; **Fig 1B**). Then, Tree SHAP explanation [20] is used to assess the contribution of inputs to representations (**Materials & Methods**; **Fig 1B**). As such, XA4C outputs SHAP values for input gene expression individually and quantifies the contribution of each input to each representation (i.e., latent variable). Using these SHAP values, XA4C further quantifies the Critical index of a gene by averaging the absolute values of its SHAP value to all latent variable via all the samples (**Materials & Methods**). By ordering these genes based on their Critical indexes, XA4C achieves a list of *Critical genes* above a user-specified cutoff, e.g., top 1% (**Materials & Methods**; **Fig 1C**). As downstream applications of XA4C, the genes together with their Critical indexes may be used for pathway enrichment analysis and connectivity analysis (**Materials & Methods**; **Fig 1D, E**).

**Fig 1.**
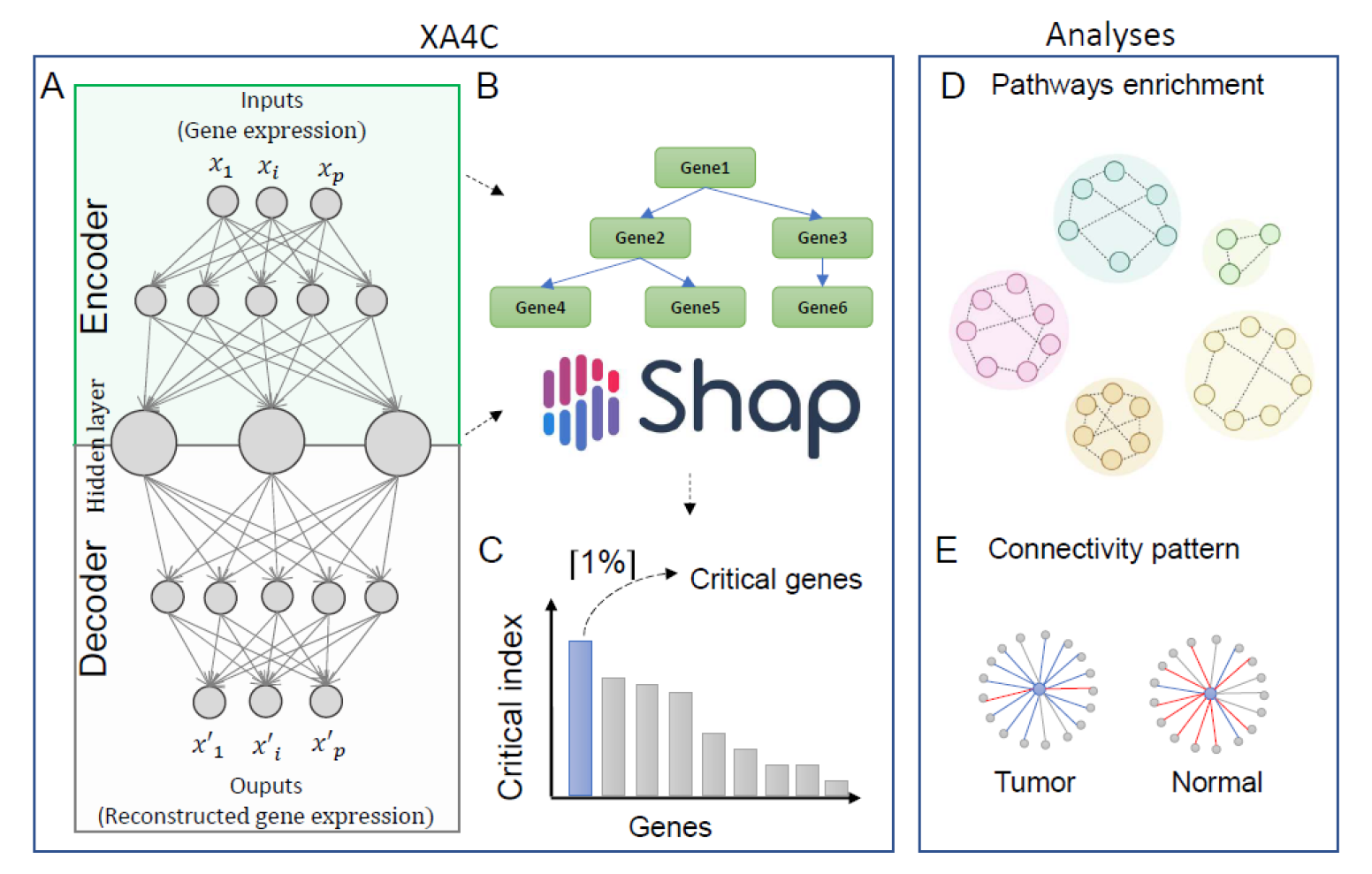
The XA4C model and potential downstream analysis. (A) An autoencoder is constructed to learn representations (i.e., latent variables) of input gene expression profiles. (B) XGBoost and TreeSHAP are utilized to evaluate SHAP values and Critical indexes for all genes. (C) Critical genes are the ones with the top 1 % Critical indexes. (D) KEGG pathway enrichment identifies sensible pathways overrepresented by prioritized genes with SHAP values. (E) Connectivity analysis discloses interaction patterns among genes centered by Critical genes in pathways.

### Whole-transcriptome AEs with high compression ratio reconstructed with high accuracy

First, we established whole transcriptome AEs based on curated 15,000 genes (**Materials & Methods**). Second, we evaluated reconstruction performances for AEs by calculating R^2^ from testing samples (**Table 1**). It is observed that AEs with 32 latent variables conducted decent reconstruction with R^2^ values from the testing dataset varying from 0.42 - 0.69, which are comparable to AEs with 512 latent variables (R^2^ values range from 0.50 - 0.72). However, the compression ratio of AEs with 32 latent variables (468≈15,000/32) is 16 times of AEs with 512 nodes (29≈15,000/512), which indicates latent variables from the former compress way more information. Therefore, for the whole transcriptome analysis of XA4C, we used AEs with 32 latent variables nodes for the six cancers. We also set up this as the default in XA4C.

**Table 1.** Test R^2^ for whole transcriptome autoencoders. L is the number of layers and H is the number of latent variables.

| Cancer | Primary Tissue | Sample size | Number of genes in AE inputs | Test $R^2$ (L=5, H=32) | Test $R^2$ (L=1, H=512) |
| --- | --- | --- | --- | --- | --- |
| BRCA | Breast | 1048 | 15,488 | 0.50 | 0.50 |
| COAD | Colon Adenocarcinoma | 389 | 14,211 | 0.47 | 0.50 |
| KIRC | Kidney | 481 | 14,459 | 0.64 | 0.62 |
| LUAD | Bronchus and lung | 485 | 15,341 | 0.42 | 0.53 |
| PRAD | Prostate gland | 459 | 14,843 | 0.52 | 0.53 |
| THCA | Thyroid gland | 451 | 14,500 | 0.69 | 0.72 |

### Whole-transcriptome Critical genes and pan-cancer pathways

We calculated Critical indexes for the ∼15,000 genes (**Supplementary Table S1**) and illustrated the top 20 in **Fig 2A**. It is observed that the maximum absolute values for Critical indexes range from 0.03 to 0.06, and they decrease quickly. The overall distribution of the Critical indexes all genes and Critical genes in six cancers are in **Fig 2B**.

**Fig 2.**
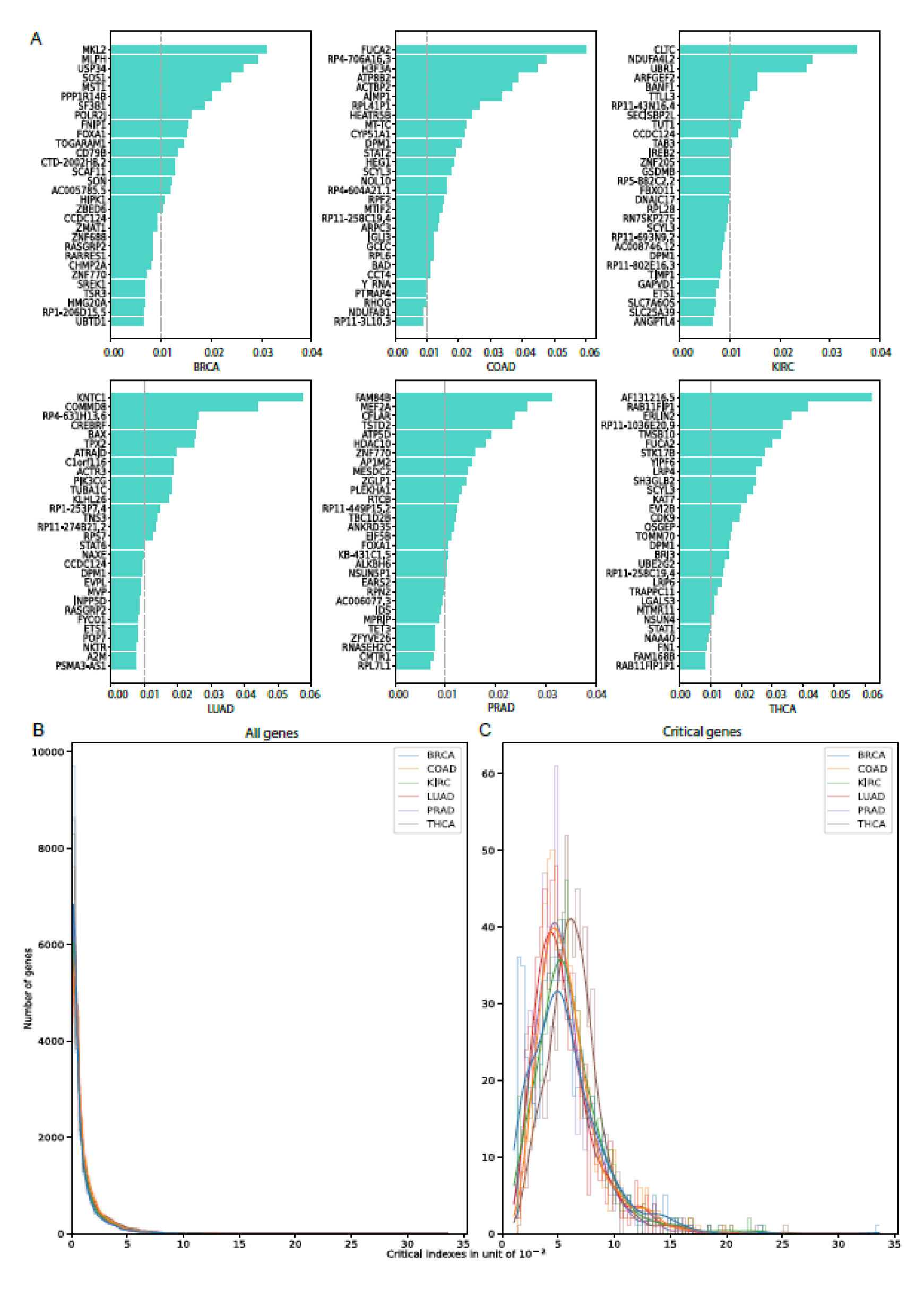
Whole transcriptome Critical indexes of genes in six cancers. (A) Genes with the largest 20 Critical indexes summarized among all latent variables and averaged across samples. (B) Distribution of whole transcriptome Critical indexes for all genes. (C) Distribution of whole transcriptome Critical indexes for Critical genes.

We further conducted pathway over-representation analysis on genes with non-zero Critical indexes to reveal sensible pathways underlying cancers (**Materials & Methods**). We found a handful of pathways that mediate crucial roles in multiple cancers (**Fig 3A; Supplementary Table S2**). Notably, oxidative phosphorylation (OXPHOS) was enriched for five cancers (BRCA, COAD, KIRC, LUAD and THCA), and growing evidence indicate it was an active metabolic pathway in many cancers [25]. Higher expression of OXPHOS genes predicts improved survival in some cancers [26], but also confers chemotherapy resistance in others [27, 28]. Many recent studies proposed to treat this pathway as an emerging target for cancer therapy [26-28]. Another interesting pathway, glutathione metabolism, has been found in two cancers (KIRC and THCA). Glutathione (GSH) is an important antioxidant that maintains cellular redox homeostasis and detoxifies carcinogens [29, 30]. However, GSH metabolism can also play a pathogenic role in cancer by conferring therapeutic resistance and promoting tumor progression [30, 31]. There are novel therapies that target the GSH antioxidant system in tumors to increase treatment response and decrease drug resistance [30]. Furthermore, amino sugar and nucleotide sugar metabolism pathway has been identified LUAD, and studies showed that arresting pathways of carbohydrate metabolism such as central carbon metabolism in cancer, aerobic glycolysis, and amino sugar and nucleotide sugar metabolism can introduce apoptosis in breast cancer [32]. Other well-known cancer pathways, such as apoptosis, p53 signalling pathways, have also been discovered by Critical index-directed enrichment analysis.

**Fig 3.**
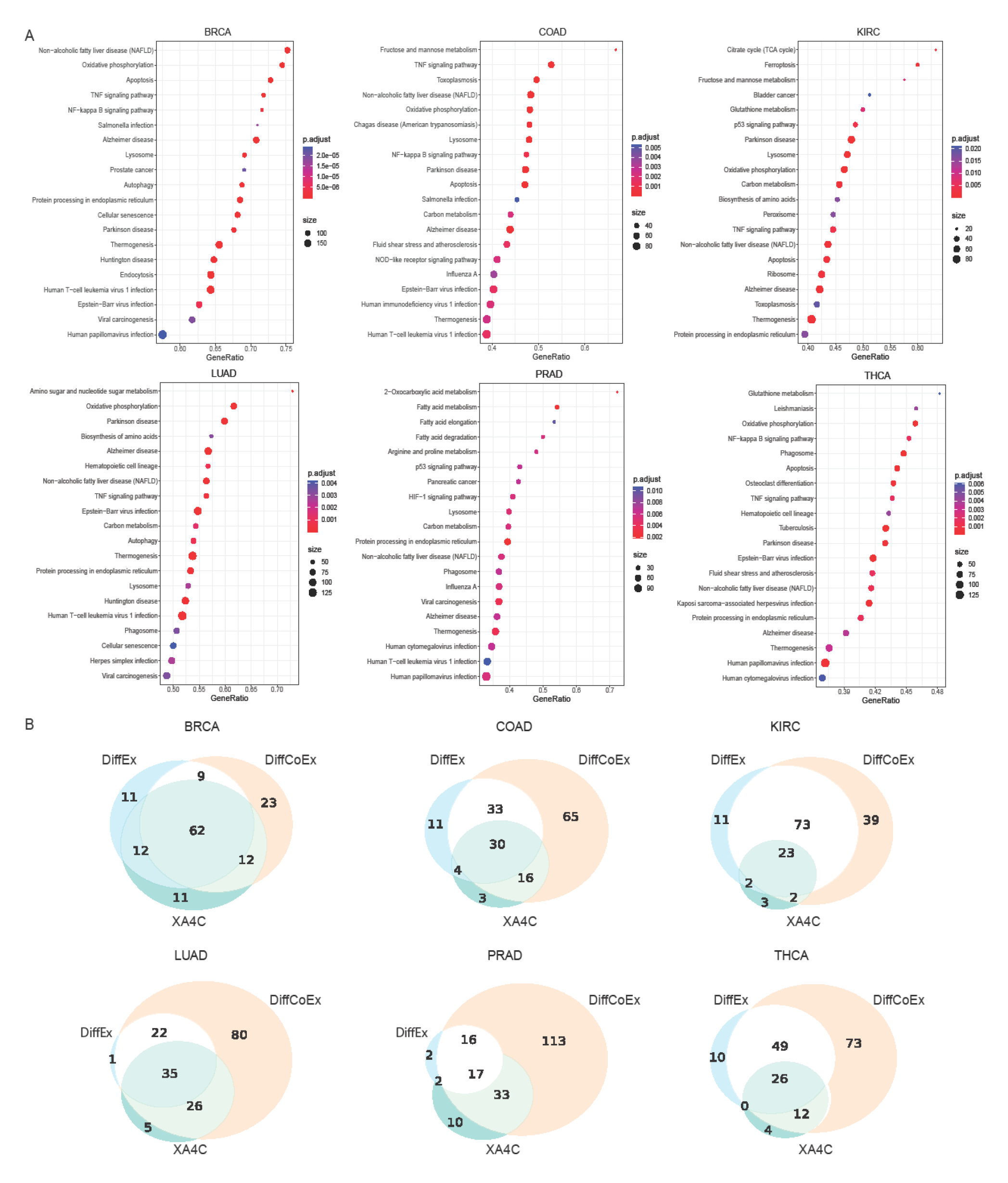
Pathway enrichment of whole-transcriptome genes. (A) Top 20 KEGG pathways enriched by genes with non-zero Critical indexes. (B) Comparison of pathways enrichment of genes prioritized by XA4C, DiffEx analysis and DiffCoEx analysis.

As comparisons, we also performed differential expression (DiffEx) analysis [33] and differential co-expression (DiffCoEx) analysis [12] (**Materials & Methods**). Overall, 31%∼83% of XA4C pathways are shared with DiffEx, and the percentage increased to 76%∼91% for DiffCoEx (**Fig 3B**). Our results showed that XA4C’s ability of identifying pathways has a larger overlap with the network-based approach DiffCoEx than the marginal effect-based approach DiffEx. It may because AE can handle nonlinear relationships among features, whereas traditional DiffEx analysis couldn’t utilize gene network information.

### Overview of within-pathway Critical genes

To further explore the Critical genes within known pathways, we applied AEs to expressions of genes within individual pathways. From the KEGG [34], we downloaded 334 pathways whose numbers of genes varies from dozens to hundreds (**Supplementary Table S3**). For each pathway of each cancer, we constructed an AE, with a small number of latent variables (H=8) and few layers (number of layers L= 3 or 2). We set L=2 when the number of genes in a pathway is less than 100, or L=3 otherwise. Pathway AEs testing R^2^ values have noteworthily high values with mean values above 0.6 for all six cancers (**Fig 4A**). It is also noted that the mean testing R^2^ values from KIRC, PRAD, and THCA pathway AEs are larger than BRCA, COAD and LUAD, which is consistent with the AEs performance of whole-transcriptome analysis. The distributions of Critical indexes for genes from 334 pathways are similar to whole-transcriptome results (**Fig S1A**) and pathway Critical genes (the largest 1%) mean values also approximate transcriptome Critical genes (**Fig S1B**).

**Fig 4.**
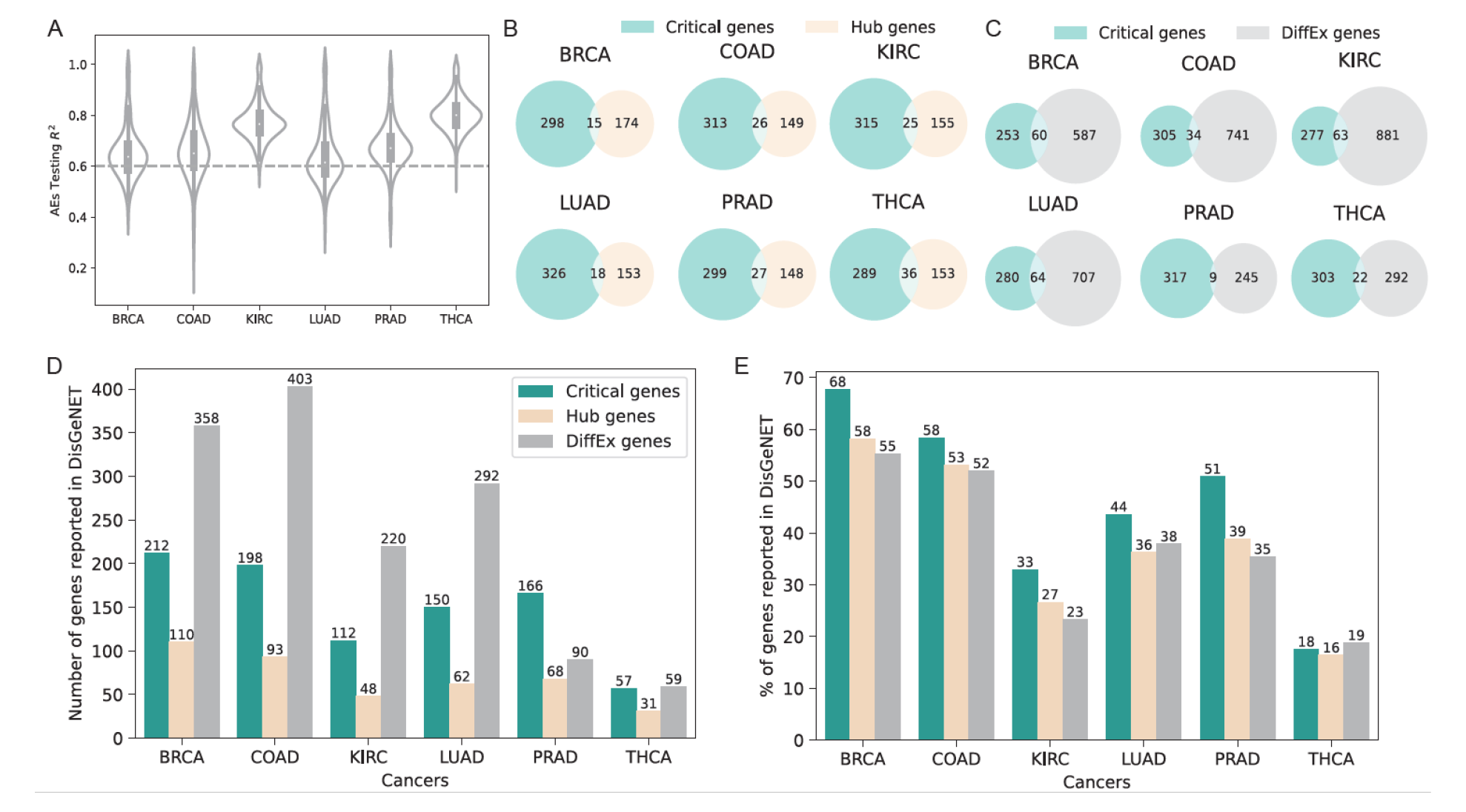
Generation and analysis of within-pathway Critical genes. (A) Distribution of R^2^ (in testing samples) of pathway AEs in six cancers. (B) Overlaps between Critical genes and Hub genes (identified by WGCNA). (C) Overlaps between Critical genes and DiffEx genes. (D) Numbers of Critical, Hub, and DiffEx genes validated by DisGeNET. (E) Percentage of Critical, Hub, and DiffEx genes validated by DisGeNET.

### The overlaps between Critical genes and Hub or DiffEx genes are poor

To reveal whether Critical genes indeed make differences in practice, we compare Critical genes to Hub genes defined by Weighted Correlation Network Analysis (WGCNA) [13] as well as DiffEx genes generated by DESeq2 [33] (**Materials & Methods**). As WGCNA outputs 1 Hub gene per pathway and our 1% Critical index cut-off in the pathway is on average approximately 1 or 2 genes, this analysis yields comparable number of genes. We found that, in all six cancers, the overlap between Critical genes and Hub genes are poor (**Fig 4B**) Similar observation is also evident for DiffEx genes (**Fig 4C**). These results show that Critical genes indeed provide a different angle for researchers to analyze expression data.

### Critical genes have higher enrichment in disease genes than Hub and DiffEx

Having learned the low overlap presented above, we then continue to learn whether the Critical genes prioritized by XA4C are indeed sensible. We then examined the DisGeNET [35], a comprehensive database for the enrichment of these three categories of genes (**Materials & Methods**). We noticed that Critical genes are highly enriched in genes with susceptibility reported in DisGeNET. Although DiffEx has the number of successfully validated genes due to its overall substantially more input candidates (**Fig 4D**), Critical gene is the winner when comparing the ratio (numbers of validated genes divided by numbers of input genes) (**Fig 4E**). Notably, the high proportions are consistent across all six cancers, indicating that Critical genes are fundamental for all cancers. Together, these results indicate that XA4C-derived Critical genes capture additional information other than marginally altered gene expression and genes with a high degree of connectivity in the protein-protein interaction network.

### Critical genes alter interaction patterns in a pan-cancer pathway

To further investigate the functions performed by Critical genes in their pathways, we calculated the Pearson correlation between the Critical genes and all the other genes in corresponding pathways. We found that Critical genes display distinct interaction patterns in the co-expression network between tumor and matched normal tissues. We specifically picked up an example in Lysine degradation (hsa00310) pathway as the Critical genes in this pathway are neither Hub nor DiffEx genes in five cancers. Evidently, the interaction patterns of Critical genes and surrounding genes are dramatically different in five cancers, which are visible by inspections and quantified by the Kolmogorov-Smirnov test (**Fig 5**). First, the intensity of correlation is notably weaker in tumor while compared to normal (lighter color in tumor and brighter color in normal). Moreover, the variance of correlations (reflected by the height of the boxplots) is substantially larger in normal tissues in all cancers except for THCA. Overall, the dramatically changed interaction patterns suggest that Critical genes involved in disease pathogenesis through interactions with other genes although themselves are not marginally differentially expressed (DiffEx genes) nor most connected with other genes (Hub genes) in the pathway.

**Fig 5.**
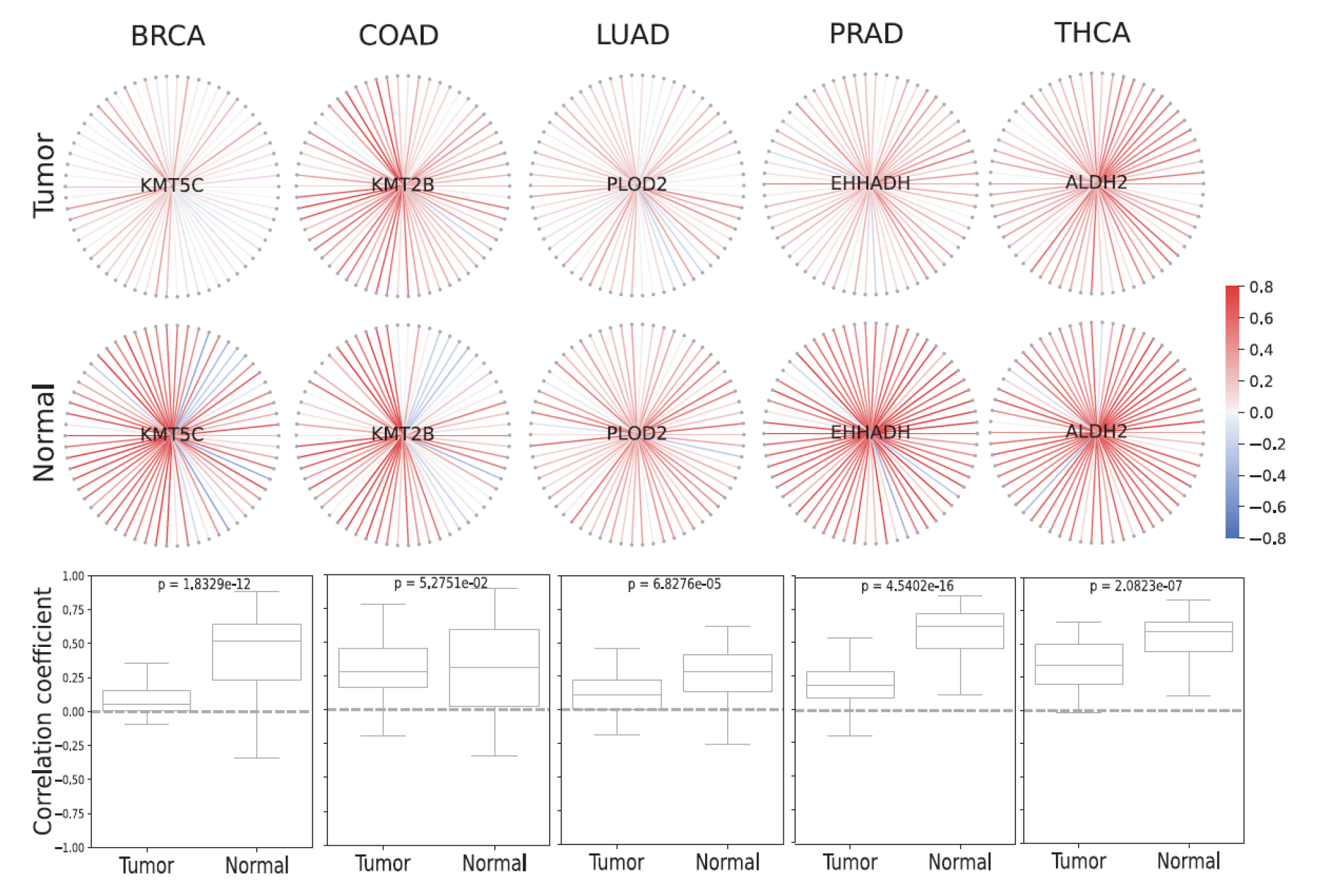
Critical genes show distinct co-expression networks in tumor and normal tissues. The Lysine degradation pathway (hsa00310) is used. Critical genes (light blue) are located at the core of the network, surrounded by additional genes from the same pathway (gray). The boundaries of Pearson’s correlation coefficients range from +0.8 (red) to -0.8 (blue). Boxplots show the distributions of two sets of correlations (tumor vs. normal) together with the P-value of the Kolmogorov-Smirnov test, with the null hypothesis being that the two samples were chosen from the same distribution. Critical genes shown in this figure are novel as they have not been identified by traditional analysis search for Hub nor DiffEx genes.

## DISCUSSION

In this work, we proposed XA4C, an XAI empowered AE tool to support explainable representation learning (RL). A notable contribution of XA4C is the definition of Critical genes, which formed the RL analogue to Hub/DiffEx genes, bringing an additional perspective in characterizing transcription data. By applying XA4C to cancer transcriptomes, XA4C generated Critical indexes and the list of Critical genes. Analyses show that Critical genes are quite different to DiffEx and Hub genes offered standard analysis based on plain representations, opening a potential landscape for in-depth analysis of complex interactions using RL. Impressively, Critical genes enjoy higher success rate in DisGeNET, a comprehensive disease gene database, indicating that Critical genes indeed play functional roles in diseases. As an example of interactions revealed by Critical genes, XA4C highlighted interesting Critical genes playing central roles in pathways by showing distinct the interaction patterns between tumor and normal tissues.

Utilizing the efforts in machine learning, researchers have developed several of tools for interpretable analysis. In particular, Hanczar and colleagues employed gradient methods to analyze the contribution of specific neurons in the network [16], which, however, does not focus on the contribution of individual input genes. Dwivedi and colleagues analyzed the effect of an input feature by differentiating the outcome by switching off the focal input feature, i.e., gene [4]. Although aiming to relieve the problem of interpretation, this method only focuses on the marginal effect of each gene, which does not employ the latest development in XAI that systematically examines complex models. Recently, Withnell et al proposed XOmiVAE [8] that also employs SHAP and AE, however it focuses more on the classification of samples, instead of prioritizing individual genes for further analysis.

We acknowledge limitations of XA4C, which could be addressed by our future work. First, we utilized fixed architectures of AEs in this study. Although we compared the performances of AEs on several different architectures and selected the optimal architectures for whole-transcriptome (L=5, H=32) and pathway AEs (L=3 or 2, H=8), we only tested a few combinations of the architecture parameters. A sophisticated way is to incorporate Bayesian Hyperparameter Optimization[36] for extensively evaluating the combinations of L and H. Second, only conventional AEs are used. It is straightforward to extend to other AEs, such as Variational AE for causal inference and Graph AE to learn causal structure, which is our planned future work. Third, TreeSHAP was utilized in this study to explain the tree model built on AEs inputs and representations. Some SHAP explainers, such as DeepSHAP [37], might be more efficient for neural network models since they omit the phase of constructing Tree models between inputs and representations. Fourth, we summarized SHAP values over samples to generate one Critical index to rank the importance of genes. This gives us general information but ignores the heterogeneity between individual samples. Our future study will focus on individuals to uncover individualized Critical genes to support precision medicine. Finally, XA4C currently supports only transcriptome data, and a natural extension will be the incorporation of additional omics data, such as proteomics and metabolomics.

## MATERIALS & METHODS

### XA4C model (I): Autoencoder (AE) architectures and parameters tuning

AE consists of two main components: an encoder and a decoder. The encoder converts the input data into a lower-dimensional representation, known as the “latent variables”. The decoder then reconstructs the original input data based on latent variables. The objective of an autoencoder is to minimize the reconstruction error between the input data and the reconstructed. From a mathematical perspective, the encoder part can be represented by the encoding function h = f(x), and the decoder can be represented by the decoding function r = g(h). Thus, the whole AE can be described by a function *r* = g(f(x)), where the output r is reconstructed as approximate as possible to the original input x. XA4C utilizes Mean Square Error 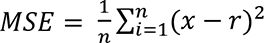 as the loss function.

By default, XA4C specifies an AE with 5 coding layers and 32 latent variables for whole transcriptome, and AEs with 3 coding layers (or 2 if the number of genes is lower than 100) and 8 latent variables for pathway gene expressions. We achieved these optimal parameters by testing configurations striking a balance between fewer AE parameters (layers, nodes) and a larger reconstruction R^2^ in testing sample. To train AE models, we partitioned tumor samples into training and testing datasets with the ratio of 8:2. Genes with median expression levels larger than 1.0 were retained as input. Raw gene expressions were transformed using Log (base 2) function, and then further rescaled to a range between 0 and 1 to match the sigmoid activation function in AE neural networks. To train AE models, we utilized the *Adam* optimizer [38] with a learning rate = 1.0 × 10^-3^ and decay = 0.8 for 500 epochs. The training stops when testing R^2^ does not increase by more than 0.02% in 10 epochs, or the maximum number of training epochs (3,000) is reached. The batch size was set as the input sample size. These AE models were implemented using PyTorch [39].

### XA4C model (II): SHapley Additive exPlanation (SHAP) framework, XGBoost and TreeSHAP

SHAP is a classical post-hoc explanatory framework to calculate the contribution of each input variable in each sample learn the explanatory effect [18]. Let *f* be the original model and *f*(*x*) the predicted values. *g*(*x*′) is the explanation model used to match the original model *f*(*x*). Note that explanation models often use simplified inputs *x*′ that map to the original inputs through a mapping function *x* = ℎ_*x*_(*x*′). XA4C incorporates **eXtreme Gradient Boosting (XGBoost)** Regressor [23] to represent the model *f* and Tree Explainer [20] as the model *g* to interpret *f* in SHAP. We choose XGBoost because models built through gradient boosting algorithm gives more importance to functional features and are less vulnerable to hyperparameters initializations [40]. Each latent variable has an XGBoost model. TreeSHAP is a SHAP method designed for the black-box tree models including Random Forest, XGBoost, and CatBoost, etc [41].For each XGBoost regression model (established between all inputs and a latent variable in the AE), XA4C carries out TreeSHAP calculation, leading to a SHAP value for each triple of [gene, latent variable, sample].

### XA4C model (III): definition of Critical indexes and Critical genes

The Critical index of a gene is the weighted average of its SHAP value contributed to all samples and all latent variables. To calculate the Critical index (*WSV_i_*) of gene *i*, XA4C first takes the mean absolute value for the gene *i* over all *n* samples, leading to the SHAP value of a gene to a latent variable:

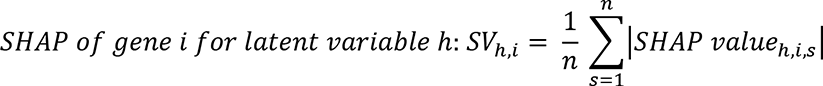

Then, XA4C summarizes SHAP values (*SV_h,i_*) across all latent variables (e.g., 32 in the default AE configuration of XA4C) and weights them by *w*_ℎ_, which is the XGBoost regressor *R*^2^ of the ℎ-th latent variable:

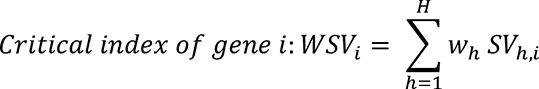

XA4C ranks all genes based on their Critical indexes and take the top ones (based on a user-specified cutoff) as Critical genes. By default, XA4C selects 1% for both whole transcriptome is and within-pathway analysis. The ceiling function ⌈ ∗⌉, i.e., the least integer greater than the actual value (e.g., ⌈ 0.2⌉ = 1, ⌈ 1.8⌉ = 2), is used when the cut-off is not an integer.

### TCGA gene expression data for six cancers

We utilized gene expression (number of genes M=56,497)) from six cancers (BRCA: breast invasive carcinoma, COAD: colon adenocarcinoma, KIRC: kidney renal clear cell carcinoma, LUAD: lung adenocarcinoma, PRAD: prostate adenocarcinoma, THCA: thyroid carcinoma) from The Cancer Genome Atlas (TCGA) [21]. The raw count data was downloaded from the TCGA data portal and then converted to Transcripts Per Million (TPM) gene expression matrices using gene lengths obtained through the BioMart package (code available in our GitHub). We then performed basic quality control to examine the overall structure of the data by PCA analysis [42] and did not find unexpected data points. In particular, we curated genes with median expression levels higher than 1.0, which reduces interference from lowly expressed genes and the number of genes decreased to around 15K. We also performed log-2 transformation to these 15K genes. When applying XA4C to cancer data, for each cancer type, one whole-transcriptome AE and 334 pathway AEs were trained independently. The architecture of the embedding network contains 32 latent variables for whole-transcriptome AE or 8 latent variables for pathway AEs. These whole-transcriptome or pathway representations and their corresponding inputs were passed through the eXtreme Gradient Boosting (XGBoost) regression model on which TreeSHAP SHAP values for inputs genes.

### Pathway over-representation analysis

Over-representation analysis (ORA) [43] is a statistical method to understand which biological pathways may be over-represented. It determines whether genes from pre-defined sets (e.g., those belonging to a specific KEGG pathway) are present more than would be expected (over-represented). The p-value can be calculated by the hypergeometric distribution: 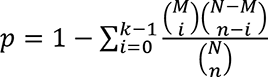, where *N* is the total number of genes in the background set, *n* represents the size of the list of genes of interest (for example the number of DiffEx or Critical genes), *M* stands for the number of genes annotated background, and *k* is the number of genes in the list annotated to the gene set. The background distribution, by default, is all the genes that have annotation, which are KEGG pathway genes in this study. We utilized the ORA analysis provide by the R package named WebGestalt [44].

### Differential expression and Differential Co-expression Analysis

Differential expression (DiffEx) analysis aims at identifying differentially expressed genes between experimental groups. We utilized DESeq2 [33], which tests for differential expression by negative binomial generalized linear models [33]. As DiffEx analysis is based on linear models, it ignores the non-linearity displayed by gene expression. Therefore, we used differential co-expression networks to identify groups (or “modules”) of differentially co-expressed genes utilizing “DiffCoEx” [12], an extension of the WGCNA[13]. DiffCoEx begins with the construction of two adjacency matrices: 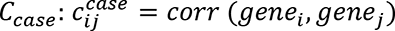 for case samples and *C_control_* similarly for control samples. DiffCoEx used the Spearman rank correlation, and a matrix of adjacency difference is then calculated:

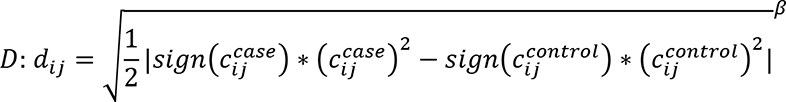

where *β* ≥ 0 is an integer tuning parameter which can be selected in multiple ways. In this study, we chose *β* ∈ [5,6,7,8,9,10] such that we achieved minimum number of modules with the largest module containing the smallest number of genes. Next, a Topological Overlap dissimilarity Matrix is calculated where smaller values of *t*_*ij*_ indicates that a pair of genes *gene_i_*, and *gene_j_* have significant correlation changes (between case and control).

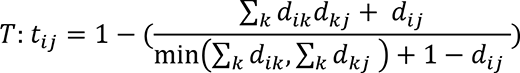

Finally, the dissimilarity matrix *T* is to identify DiffCoEx genes.

## SUPPORTING INFORMATION

**Supplementary Table S1: Critical indexes for whole transcriptome AEs in six cancers**

**Supplementary Table S2: KEGG pathways enrichment for XA4C, DiffEx and DiffCoEx**

**Supplementary Table S3: KEGG pathways and numbers of genes in pathways**

**Supplementary Fig S1.**
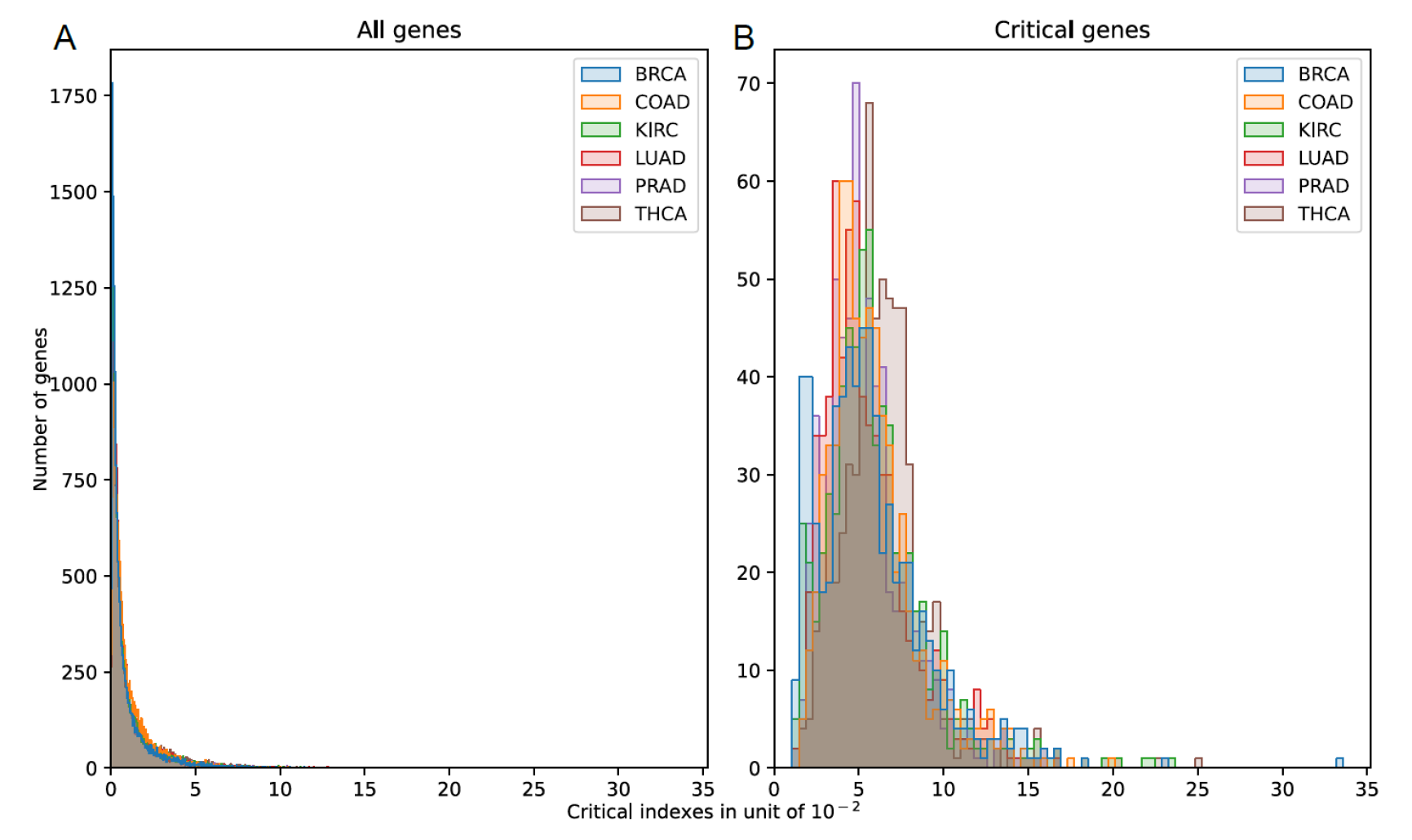
Pathway Critical indexes of genes in six cancers. (A) Distribution of pathway Critical indexes for all genes in the corresponding pathways. (B) Distribution of pathway Critical indexes for Critical genes in the corresponding pathways.

## AVAILABILITY OF DATA AND MATERIALS

XA4C is publicly available in our GitHub: https://github.com/QingrunZhangLab/XA4C

TCGA: https://portal.gdc.cancer.gov/

DisGeNET: https://www.disgenet.org/

DESeq2: http://www.bioconductor.org/packages/release/bioc/html/DESeq2.html

WGCNA: https://cran.r-project.org/web/packages/WGCNA/index.html

## Author contributions

Conceptualization: Q.Z.

Data curation: Q.L., Y.Y., P.K.

Formal analysis: Q.L., Q.Z.

Funding Acquisition: Q.Z., W.L.

Investigation: Q.L., Q.Z.

Methodology: Q.L., Q.Z.

Resources: W.L., Q.Z.

Software: Q.L., P.K.

Supervision: W.L., Q.Z.

Visualization: Q.L., T.L.

Writing—original draft Preparation: Q.L., Q.Z., with contributions from all co-authors

Writing—review & editing: Q.L., and Q.Z.

