## Supplementary files for "XA4C: eXplainable representation learning via Autoencoders revealing Critical genes"

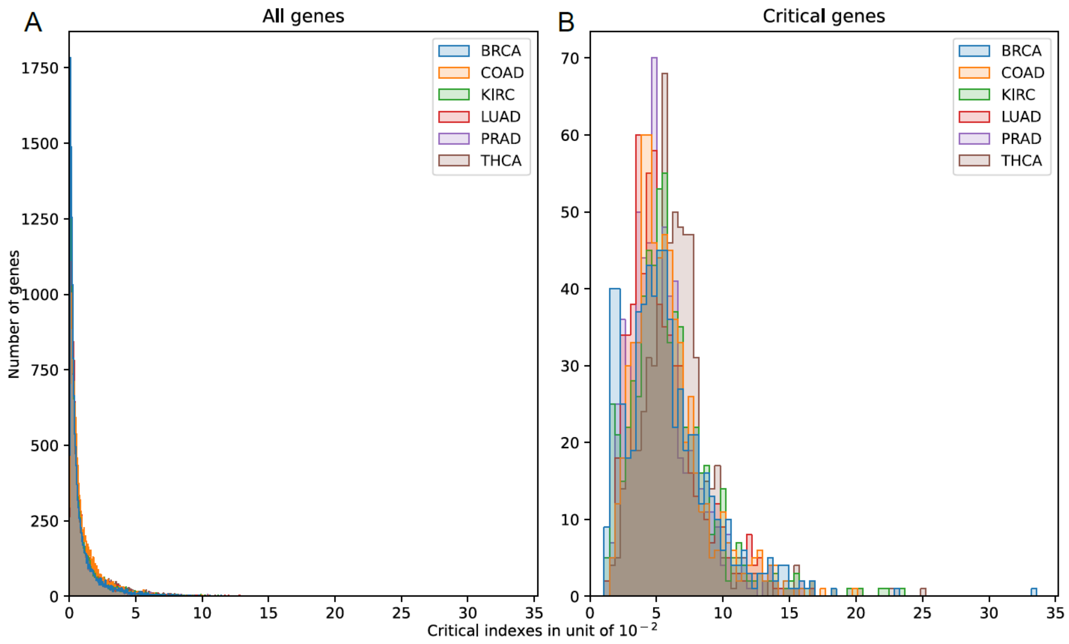


**Supplementary Fig S1. Pathway Critical indexes of genes in six cancers.** (A) Distribution of pathway Critical indexes for all genes in the corresponding pathways. (B) Distribution of pathway Critical indexes for Critical genes in the corresponding pathways.
